# Identification of root rot resistance QTLs in pea using *Fusarium solani* f. sp. *pisi*-responsive differentially expressed genes

**DOI:** 10.1101/2020.11.13.382077

**Authors:** Bruce A. Williamson-Benavides, Richard Sharpe, Grant Nelson, Eliane T. Bodah, Lyndon D. Porter, Amit Dhingra

## Abstract

*Pisum sativum* (pea) yields have declined significantly over the last decades, predominantly due to susceptibility to root rot diseases. One of the main causal agents of root rot is the fungus *Fusarium solani* f. sp. *pisi* (*Fsp*), leading to yield losses ranging from 15 to 60%. Determining and subsequently incorporating the genetic basis for resistance in new cultivars offers one of the best solutions to control this pathogen; however, no green-seeded pea cultivars with complete resistance to *Fsp* have been identified. To date, only partial levels of resistance to *Fsp* has been identified among pea genotypes. SNPs mined from *Fsp*-responsive differentially expressed genes (DEGs) identified in a preceding study were utilized to identify QTLs associated with *Fsp* resistance using composite interval mapping in two recombinant inbred line (RIL) populations segregating for partial root rot resistance. A total of 769 DEGs with single nucleotide polymorphisms (SNPs) were identified, and the putative SNPs were evaluated for being polymorphic across four partially resistant and four susceptible *P. sativum* genotypes. The SNPs with validated polymorphisms were used to screen two RIL populations using two phenotypic criteria: root disease severity and plant height. One QTL, *WB.Fsp-Ps* 5.1 that mapped to chromosome V explained 14.76 % of the variance with a confidence interval of 10.36 cM. The other four QTLs located on chromosomes II, III, and V, explained 5.26–8.05 % of the variance. The use of SNPs derived from *Fsp*-responsive DEGs for QTL mapping proved to be an efficient way to identify molecular markers associated with *Fsp* resistance in pea. These QTLs are potential candidates for marker-assisted selection and gene pyramiding to obtain high levels of partial resistance in pea cultivars to combat root rot caused by *Fsp*.

## Introduction

Pea (*Pisum sativum* L.; Family Fabaceae) is an important cool-season, self-pollinating annual diploid crop. A number of cultivars within the species cater to different consumption markets. Green peas, and dry yellow and green peas are sold as food in the fresh and dry markets, respectively, while purple-seeded lines are used for forage and green manure (Miller et al., 2005). Due to its high protein content (20–30%) and overall high nutritional value, pea has become a major contributor to the plant-derived protein market (do Carmo et al., 2016; Peng et al., 2016; Xiong et al., 2018; Wei et al., 2020). The shift to plant-based protein is an environmentally sustainable alternative to animal-based protein because the latter contributes significantly to greenhouse gas emissions (Stehfest et al., 2009). Furthermore, studies have shown that dietary proteins in peas are of great benefit to human health and wellness (Reddy and Yang, 2011; Kudre et al., 2013; Dahiya et al., 2015). Consequently, the pea protein market is projected to reach $34.8 million in 2020 (Grand View Research, 2015; World Health Organization, 2015; Pietrysiak et al., 2018).

The profitable production of pea is threatened by soilborne diseases. These diseases are commonly referred to as the pea root rot complex (PRRC) and are caused by a single or combination of pathogens, including *Aphanomyces euteiches*, Fusarium spp, *Mycosphaerella pinodes*, Pythium spp., and *Rhizoctonia solani* (Xue, 2003; Kumari and Katoch, 2020). One of the predominant causal agents of PRRC is the fungus *Fusarium solani* f. sp. *pisi* (*Fsp*). *Fsp* occurs in most pea fields throughout the world, and the yields of *P. sativum* cultivars can be reduced up to 15-62% by this pathogen (Seaman, 1976; Grünwald et al., 2003). *Fsp* infects pea seeds during germination, with symptoms of root rot beginning at or near the cotyledon-hypocotyl junction and progressing under the soil and upper region of the taproot (JM, Kraft and Pfleger, 2006). Round or irregular light brown lesions that progress to dark black lesions on below-ground stems have also been reported, along with stunting and death (Jung et al., 1999). *Fsp* can survive in the soil for more than one season and conditions that decrease root growth, such as soil compaction, extreme temperatures, and moisture levels, can increase *Fusarium*-mediated root damage (Shaykh et al., 1977; JM, Kraft and Pfleger, 2006).

The development of pea cultivars with root rot resistance has been considered the best long-term management option among the many root rot control strategies (Conner et al., 2014; Bodah et al., 2016; Wang et al., 2018). However, breeding for *Fsp* resistance is challenging, since resistance to *Fsp* is a quantitative trait (Mukankusi et al., 2011; Román-Avilés et al., 2011; Bodah et al., 2016). Furthermore, routine screening for resistance has proven to be time-consuming, expensive, and highly influenced by the environment (Bodah et al., 2016).

Marker-assisted selection (MAS) can help expedite the selection of putative *Fsp* resistant progeny without the need for expensive phenotyping. Several efforts have been made to develop molecular markers associated with resistance to *Fsp* root rot in pea (Feng et al., 2011; Coyne et al., 2015, 2019). However, these studies used a limited number of DNA markers and some of the QTLs identified require further fine mapping due to large confidence intervals (16.8-28.5cM).

In a preceding time-course transcriptome study, we utilized a combined genetic and RNAseq approach to identify *Fsp*-responsive differentially expressed genes among four partially resistant and four susceptible genotypes (Williamson-Benavides et al., 2020). These genotypes were selected for their contrasting root severity index phenotype (Bodah et al., 2016). Genes involved in secretion and exocytosis, anthocyanin biosynthesis pathway genes, and a previously-described pathogenesis-related (PR) gene DRR230 were observed to be overexpressed in partially resistant genotypes (Chiang and Hadwiger, 1991; Williamson-Benavides et al., 2020). Since the use of single nucleotide polymorphisms (SNPs) can help to refine genetic mapping studies due to their high abundance in the genome (Deulvot et al., 2010), SNPs mined from differentially expressed genes (DEGs) were utilized to identify QTLs associated with *Fsp* resistance using composite interval mapping in two recombinant inbred line (RIL) populations segregating for root rot resistance as observed in greenhouse evaluations.

## Materials and Methods

### Plant material

The parental plant material used in this study was the same as described in a preceding study (Williamson-Benavides et al., 2020). Briefly, four genotypes with partial resistance to *Fsp*—00-5001, 00-5003, 00-5004, and 00-5007— and four susceptible genotypes— ‘Aragorn’, ‘Banner’, ‘Bolero’, and ‘DSP’— were selected based on their disease resistance to *Fsp* (Table 1). These genotypes were previously classified as either partially resistant or susceptible based on phenotyping root disease severity index (RDS), plant height, shoot dry weight, and root dry weight after *Fsp* challenge (Bodah et al., 2016). The 5000 series pea breeding lines were found to be the most resistant lines among the white-flowered pea lines. The susceptible genotypes are among the most frequently used commercial pea varieties in the United States (Table 1).

**Table 1.** Selected green-seeded pea genotypes for SNP genotyping (Adapted from Williamson-Benavides et al., 2020). The table shows seed source, *Fsp* tolerance level, other disease resistance, 100 seed weight, leaf type, and market class per pea genotype.

| Genotype | Source <sup>a</sup> | *Fsp* resistance level <sup>b</sup> | Other disease resistance <sup>c</sup> | 100 seed weight | Leaf type <sup>d</sup> | Market Class |
| --- | --- | --- | --- | --- | --- | --- |
| 00-5001 | USDA-ARS VFCRU | * | Fop races 1, 2 and 5 | 22.7 | af | Green fresh |
| 00-5003 | USDA-ARS VFCRU | * | Fop races 1, 2 and 5 | 15.9 | af | Green fresh |
| 00-5004 | USDA-ARS VFCRU | * | Fop races 1, 2 and 5 | 20.8 | af | Green fresh |
| 00-5007 | USDA-ARS VFCRU | * | Fop races 1, 2 and 5 | 22.2 | P | Green fresh |
| Aragorn' | ProGene | *** | Fop races 1, 2; PSBMV | 19.5 | af | Green dry |
| Banner' | ProGene | *** | Fop race 2, PM | 18.7 | af | Green dry |
| Bolero' | AsGrow | **** | Fop race 1, PM, Pythium, EMV | 20.12 | P | Green fresh |
| DSP' | Canner Seed | *** | - | 20.9 | P | Green fresh |

<sup>a</sup>AsGrow = AsGrow Seed Co., San Juan Bautista, CA; Canner Seed=Canner Seed Co., Idaho Falls, ID; ProGene = ProGene LLC Plant Research, Othello, WA; USDA-ARS VFCRU=USDA-ARS, Vegetable and Forage Crops Research Unit, Prosser, WA.
<sup>b</sup>*Fsp* resistance = *Fsp* resistance resulted from phenotyping, from most resistance (\*) to most susceptible (\*\*\*\*) (Bodah et al. 2016)
<sup>c</sup>Fop= *Fusarium oxysporum*; PSBMV= Pea Seed-Borne Mosaic Virus, PM= Powdery Mildew; EMV= Enation Mosaic Virus
<sup>d</sup>af (Afila) = semi-leafless; P (Perfection) = normal leaf type

The 00-5001, 00-5003, 00-5004, and 00-5007 pea breeding lines were developed by Porter et al., (2014) via single-seed descent at USDA–ARS, Prosser, WA. The parentage of 00-5001 is PH14-119/M7477// Coquette/3/86-2197/74-410-2 (Kraft, 1989; USDA–ARS NGRP, 2020). The parentage of 00-5003 is 69PH42-691004/Recette//Popet/3/PH14-119/ DL-1/3/B563-429-2/PI257593//DSP TAC (USDA–ARS NGRP, 2020). The parentage of 00-5004 is 79-2022/ICI 1203-1//Menlo/3/PI189171/ DL-2//75-786 (Kraft and Tuck, 1986; USDA–ARS NGRP, 2020). The parentage of 00-5007 is 00-5005/ 00-5006. 00-5005 parentage is B669-87-0/M7477//Blixt B5119/3/ 00-5001/74SN5/3/PH14-119/DL-1//74SN3/Recette/5/ FR-725 (Kraft and Giles, 1976; USDA–ARS NGRP, 2020). The parentage of 00-5006 is 00-5003/00-5004.

Two F_7_-derived recombinant inbred line (RIL) populations with 190 individuals each derived from the crosses ‘Aragorn’ × 00-5001 (Population I) and ‘Banner’ × 00-5007 (Population II) were developed by single-seed descent and maintained at ProGene LLC Plant Research, Othello, WA.

### Disease challenge and greenhouse evaluations of disease

For the two populations, a total of 190 individuals with four replicates each were challenged with three *Fsp* isolates: Fs 02, Fs 07, and Fs 09. These isolates were obtained from infected pea roots collected in the Palouse Region of Washington and Idaho by Dr. Lyndon Porter, USDA-ARS Vegetable and Forage Crops Research Unit, Prosser, WA (United States). The three isolates were single-spored and were identified based on the partial translation elongation factor 1-alpha sequences (Geiser et al., 2004). The pathogenicity of each *Fsp* isolate to pea was also confirmed (Bodah et al., 2016). The three isolates were grown on pentachloronitrobenzene (PCNB) selective media for six days (Nash and Snyder, 1962). Cultures were transferred to KERR’s media (Kerr, 1963) and incubated on a shaker at 120 rpm under continuous light for six days at 23 to 25°C. The spore concentration of each isolate was determined with a hemocytometer and diluted to 1×10^6^ spores/ml of water. A spore suspension inoculum was created with equal parts by volume from each of the three isolates.

RIL seeds were surface sterilized in a 0.6% sodium hypochlorite solution and rinsed in sterile distilled H_2_O. The seeds were then soaked for 16 hours in the *Fsp* spore suspension as described previously (Bodah et al., 2016). After the challenge with the spore suspension, seeds were planted in a completely randomized design in plastic planter cones (Conetainer, 0.25 L volume, Stuewe and Sons Inc.) filled with a standard perlite medium in a greenhouse at Crites Seed Inc. (Moscow, ID). Plants were irrigated as needed, generally every 24-36 hours, and the perlite was watered to saturation at 100% field capacity. A 12-hour photoperiod was maintained using 400-watt metal halide lamps for supplemental light. Plants were grown at temperatures ranging between 21-27°C during the day and 12-18°C at night.

Quantitative evaluation of RDS and plant height were recorded 21 days after planting. RDS was evaluated on a visual scale from 0 to 6, in which 0= no diseases symptoms; 1 = small hypocotyl lesions; 2 = lesions coalescing around epicotyls and hypocotyls; 3 = lesions starting to spread into the root system with some root tips infected; 4 = epicotyl, hypocotyl and root system almost completely infected and limited white, uninfected tissue visible; 5 = completely infected root; and 6 = plant failed to emerge (Bodah et al., 2016). Plant height is a reliable indication of resistance to *Fsp* and height showed the highest negative correlation among all growth parameters related to RDS (Bodah et al., 2016). RDS and height data across the four replicates were averaged for each RIL for further analyses. Infected root tissue from three inoculated plants was taken at random to verify the presence of *Fsp* in infected tissue. The root tissue was surface sterilized and plated onto PCNB. Culture morphology and growth were observed under a microscope and compared with the original cultures to verify the presence of *Fsp* in the infected tissue.

Broad-sense heritability was estimated with this equation Va/[Va + Ve], where Va represented the genetic variance, Ve the environmental variance.

### DNA extraction

Leaf tissue was freeze-dried in a lyophilizer. Leaf tissue samples included the eight white-flowered parental genotypes —00-5001, 00-5003, 00-5004, and 00-5007 ‘Aragorn’, ‘Banner’, ‘Bolero’, and ‘DSP’, as well as the 380 RILs from Population I and Population II. DNA was extracted with the BioSprint 96 DNA Plant kit (Qiagen, Mainz, Germany). A Nanodrop ND-8000 Spectrophotometer (ThermoFisher, MA, USA) was used to quantify the extracted DNA.

### SNP mining in DEGs, SNP validation and RIL genotyping

A time-course RNAseq analysis, performed on sets of partially resistant and susceptible genotypes after *Fsp* challenge resulted in the identification of 42,905 differentially expressed contigs (DECs) (Williamson-Benavides et al., 2020). SeqMan Pro (DNASTAR, WI, USA) and custom scripts were utilized to identify single nucleotide polymorphisms (SNPs) within the set of 42,905 DECs.

The Assay Design Suite software (Agena Bioscience, CA, USA) and the SNP report generated by SeqMan Pro were used to generate two sets of primers for amplifying SNP containing regions (Table S1). The high-throughput MassArray Technology was used to validate the SNPs. Genotype calling was done from the samples deposited on the chips with the MassARRAY RT v 3.0.0.4 software (Agena Bioscience, CA, USA). Results were analyzed with the MassARRAY Typer v 3.4 software (Agena Bioscience, CA, USA). SNPs were validated across eight pea genotypes ‘Aragorn’, ‘Banner’, ‘DSP’, ‘Bolero’, 00-5001, 00-5003, 00-5004, 00-5007, which included the four parents of the two segregating populations. Each SNP was screened twice for each individual. SNPs confirmed to be polymorphic between ‘Aragorn’ × 00-5001, and ‘Banner’ × 00-5007 were used -for genotyping of 190 RILs each from Population I and Population II.

### Physical map and QTL detection

The physical location of the SNPs used in this study was determined using the pea genome (Kreplak et al., 2019). The SNP marker sequence was aligned via BLAST against the complete *P. sativum* genome in URGI BLAST (https://urgi.versailles.inra.fr/blast/).

RDS and height averages for each RIL were used to map the QTLs associated with resistance to *Fsp*. QTLs were detected with the composite interval mapping (CIM) function of the R statistical software version 3.0.2 (R core team, Vienna, Austria). CIM default settings were used. The Kosambi map function was applied to impute missing marker genotype data. QTLs were considered significant above the threshold LOD score 3.0. QTLs were named with the prefix *WB.Fsp-Ps* followed by the chromosome number and the QTL number within the chromosome.

### Functional annotation of QTLs associated with Fsp resistance in pea

The QTLs *WB.Fsp-Ps* 5.1, *WB.Fsp-Ps* 5.2, *WB.Fsp-Ps* 5.3, *WB.Fsp-Ps* 2.1, and *WB.Fsp-Ps* 3.1 were annotated using the functional annotation and gene ontology (GO) data generated in a preceding study (Williamson-Benavides et al., 2020). QTL annotation provided the identity of genes and DECs located within the selected genomic regions. The confidence intervals for each of the QTLs were taken into account to identify the genomic sequence of each QTL from the pea genome (Kreplak et al., 2019). The transcriptome data were aligned via BLAST against the QTL sequence regions in CLC Bio Genomics Workbench 6.0.1 (CLC Bio, Aarhus, Denmark).

## Results

### Disease challenge and greenhouse phenotypic evaluation

The quantitative evaluation of RDS and plant height was averaged per RIL across the four replicates as there was no significant difference between the replicates (p<0.05). Frequency histograms for both traits per population are presented in Figure 1. The phenotypic means for the parents for Population I were Aragorn-RDS=4.5; Aragorn-Height=7.0; 00-5001-RDS=2.5; and 00-5001-Height=10.5. The phenotypic means for the parents for Population II were Banner-RDS=3.25; Banner-Height=13.0; 00-5007-RDS=2.25; and 00-5007-Height=10.0. The two RIL populations displayed transgressive segregation for both increased susceptibility and resistance over the two parental lines as measured by RDS and height traits (Fig. 1).

**Figure 1.**
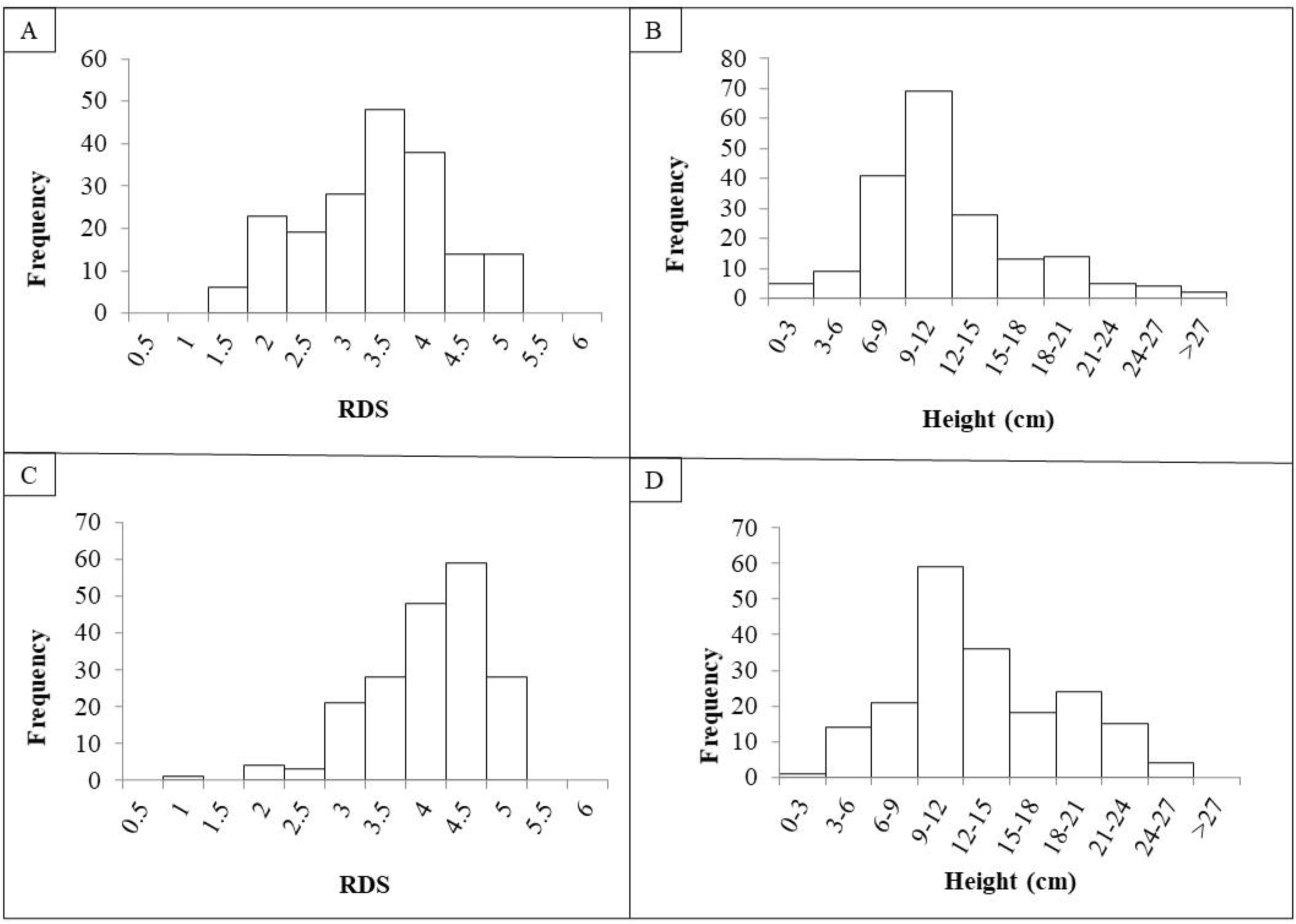
Frequency histograms of root disease severity (RDS) and plant height of recombinant lines (RILs) after challenge with *Fusarium solani* f. sp. *pisi.* RILs were derived from crosses ‘Aragorn’ × 00-5001 (A, B) and ‘Banner’ × 00-5007 (C, D).

Based on the Shapiro–Wilk test, in Population I, data were not normally distributed for RDS (W(189)= 0.95, p<0.01) or for height (W(189)= 0.91, p<0.01). Similarly in Population II, data were not normally distributed for RDS (W(189)= 0.90, p<0.01) or for height (W(189)= 0.97, p<0.01). A significant negative correlation was found between the RDS and height values for Population I (r (188)=−2.76, p<0.01) and Population II (r (188)=−4.38, p<0.01). For Population I, broad sense heritability was 49.77% and 70.44% for RDS and height, respectively. For Population II, broad sense heritability was 43.12% and 83.39% for RDS and height, respectively.

### SNP screening, SNP validation and RIL genotyping

SeqMan Pro (DNASTAR, WI, USA) identified a total of 769 SNPs across DECs (Table S2). The predicted SNPs were validated in the ‘Aragorn’, ‘Banner’, ‘DSP’, ‘Bolero’, 00-5001, 00-5003, 00-5004, and 00-5007 pea genotypes (Table S3). A total of 118 SNPs were confirmed for cultivars DSP and Bolero while 256 SNPs were confirmed for 5007 and Banner (Table 2). SNPs confirmed to be polymorphic between ‘Aragorn’ × 00-5001 (219 SNPs) and ‘Banner’ × 00-5007 (256 SNPs) were used to screen 190 individuals each of RIL Populations I and II, respectively. The screening results of 190 individuals each for both Population I and II are summarized in Tables S4 and S5, respectively.

**Table 2.** Number of SNPs across eight Pisum sativum genotypes from a total of 769 predicted SNPs.

| Genotype | Aragorn | Banner | Bolero | DSP | 5001 | 5003 | 5004 | 5007 |
| --- | --- | --- | --- | --- | --- | --- | --- | --- |
| Aragorn | - | 162 | 234 | 229 | 219 | 225 | 214 | 227 |
| Banner | 162 | - | 224 | 239 | 252 | 219 | 231 | 256 |
| Bolero | 234 | 224 | - | 118 | 191 | 220 | 191 | 213 |
| DSP | 229 | 239 | 118 | - | 206 | 223 | 177 | 218 |
| 5001 | 219 | 252 | 191 | 206 | - | 157 | 166 | 149 |
| 5003 | 225 | 219 | 220 | 223 | 157 | - | 200 | 181 |
| 5004 | 214 | 231 | 191 | 177 | 166 | 200 | - | 208 |
| 5007 | 227 | 256 | 213 | 218 | 149 | 181 | 208 | - |

### Physical map and QTL detection

The physical genomic location of all the SNPs used in this study was determined using the pea genome (Kreplak et al., 2019) (Table S6). Chromosome 1, 2, 3, 4, 5, 6, and 7 registered a total of 92, 86, 84, 122, 118, 78, 139 SNPs, respectively (Table S6). A total of 47 SNP markers were identified in 42 scaffolds that had not been assigned to any of the seven chromosomes of pea (Table S6). Three of the predicted SNP markers were not localized on the pea genome.

Prior to QTL mapping for Population I and II, a quality assessment of the genotypic data was performed. Individuals and markers with more than 80% of missing data were omitted in each database. Markers with distorted segregation patterns were also removed from the data for QTL analysis. A total of 190 RILs and 100 markers were used for QTL analysis of Population I (Table S7). A total of 182 individuals and 154 markers were used for QTL analysis of Population II (Table S8). Means per RIL for RDS and height were used to map the QTLs associated with resistance to *Fsp*. Five different QTLs were identified in the two RIL populations for RDS and height (Table 3). These QTLs explained 5.26 to 14.76% of the phenotypic variance (Table 3).

**Table 3.** Quantitative trait loci detected for resistance to *Fusarium solani* f. sp *pisi* root rot in two RIL populations using RDS and height. The RIL populations were derived from the following crosses: ‘Aragorn’ × 00-5001 (Population I) and ‘Banner’ × 00-5007 (Population II).

| Population | Scoring Phenotype | QTL Name | Chromosome | Position | Marker | LOD peak | LOD-1.5 support interval (cM) | LOD-1.5 support interval (nt) | R<sup>2</sup> |
| --- | --- | --- | --- | --- | --- | --- | --- | --- | --- |
| Population I | RDS | WB.Fsp-Ps 5.1 | 5 | 42.88 | 1_14714 | 3.68 | 40.84 – 43.98 | 236,573,488 – 254,762,537 | 8.31 |
|  | Height | WB.Fsp-Ps 5.2 | 5 | 95.39 | 7401 | 4.43 | 89.07 – 98.24 | 515,954,961 – 569,073,935 | 7.5 |
| Population II | RDS | WB.Fsp-Ps 3.1 | 3 | 9.9 | 26703 | 3.36 | 6.02 – 17.91 | 34,871,998 – 103,747,090 | 8.05 |
|  |  | WB.Fsp-Ps 5.3 | 5 | 19.23 | 3_31807 | 3.14 | 17.58 – 22.63 | 101,835,502 – 131,088,590 | 5.26 |
|  | Height | WB.Fsp-Ps 5.1 | 5 | 42.88 | 1_14714 | 6.37 | 40.83 – 51.19 | 236,515,561 – 296,527,837 | 14.76 |
|  |  | WB.Fsp-Ps 2.1 | 2 | 58.23 | 20326 | 3.17 | 56.98 – 63.40 | 330,067,516 – 367,256,590 | 5.61 |

### Functional annotation of QTLs associated with Fsp resistance in pea

The transcriptome data, generated previously (Williamson-Benavides et al., 2020) were aligned via BLAST with the 5 QTLs: *WB-Fsp-Ps* 5.1 (Table S9), *WB-Fsp-Ps* 5.2 (Table S10), *WB-Fsp-Ps* 5.3 (Table S11), *WB-Fsp-Ps* 2.1 (Table S12), and *WB-Fsp-Ps* 3.1 (Table S13). From the total set of aligned genes, 119-133 genes per QTL had previously been classified as differentially expressed (Tables S9 to S13). A total of 3-17 DEGs and 6-11 non-DEGs were also identified, predicted as having unknown function or annotated as hypothetical proteins in QTLs *WB-Fsp-Ps* 5.1, 5.2, 5.3, 2.1, and 3.1 (Tables S9 to S13).

A total of 7 DEGs associated with disease response were found in *WB-Fsp-Ps* 5.1. These genes are involved in the synthesis of lipids (acetyl-CoA carboxylase); cell signaling (C-type lectin receptor-like tyrosine-protein kinase and MAPK); nodulation (nodulation-signaling pathway 2 protein), and protein degradation (F-box/kelch-repeat). Another ten genes in *WB-Fsp-Ps* 5.1 were associated with disease resistance; however, these genes did not exhibit differential expression (Williamson-Benavides et al., 2020). These set of genes is associated with synthesis of lipids (1 gene); auxin signaling (2), ethylene synthesis (1), pectin synthesis (1), and regulation of transcription (5).

Seventeen genes found in *WB-Fsp-Ps* 5.2 were associated with disease response and also showed differential expression after *Fsp* challenge. This list included three PR (pathogenesis-related) genes (universal stress protein PHOS32-like, endochitinase PR4, protein enhanced disease resistance 2); an anthocyanin 5-aromatic acyltransferase; four receptor-like kinases; and seven TFs of the GATA, NLP8, C2H2, and scarecrow types. Another set of seventeen contigs were identified as candidate genes but did not show any differential expression. The genes on the latter list are associated with synthesis of lipids (ketoacyl-CoA synthase); three transcription factors (GLABRA and PosF21 type); an endochitinase PR4; four receptor like-kinases; and two universal stress proteins PHOS32.

Twenty-two DEGs in *WB-Fsp-Ps* 5.3 were associated with disease response mechanism. This list included drug transporters (ABC transporters); a cluster of seven F-box proteins; genes involved in cell wall biosynthesis and modification (pectinesterase/pectinesterase inhibitor and polygalacturonase); the TFIIS TF; and two PR proteins – protein enhanced disease resistance 4-like and pathogenic type III effector avirulence factor. Three more ABC transporters were found in the *WB-Fsp-Ps* 5.3, but they did not show any differential expression. Other candidate genes found in *WB-Fsp-Ps* 5.3 that did not show differential expression included an autophagy-related protein; a brassinosteroid receptor; another F-box gene; five more receptor kinases; a protein enhanced disease resistance 4-like and pathogenic type III effector avirulence factor; two TFs (CCHC(Zn) family and ERF110); and a UDP-glucuronate:xylan alpha-glucuronosyltransferase 1.

Fourteen DEGs involved in disease resistance mechanism were associated with *WB-Fsp-Ps* 2.1. These genes are known to participate in cell membrane synthesis and modification (CSC1 protein and sphingolipid transporter); PR gene response (disease resistance protein RPM1 and disease resistance protein RGA3); regulation of transcription (Myb/SANT and ninja-family protein AFP3); and cell signaling (receptor-like protein kinase 2). Three genes involved in cell wall and membrane synthesis/modification (glycerol-3-phosphate acyltransferase, sphingolipid transporter, and 3UDP-arabinopyranose mutase); and nine TFs (Myb/SANT, ninja-family protein AFP3, PLATZ transcription factor family protein) were also identified as potential candidates that contribute to the effect of *WB-Fsp-Ps* 2.1. However, this set of genes did not show differential expression after *Fsp* challenge.

Eleven DEGs associated with disease response were found within *WB-Fsp-Ps* 3.1. These genes were annotated as ethylene response sensors; polygalacturonase inhibitors; phopholipases; receptor kinases; and NDR1/HIN1-like protein 10. A set of fourteen genes were characterized as associated with disease response however they did not show differential expression. This list contains cathepsin B-like protease 2; ethylene-insensitive protein 2; F-box protein PP2-A15 isoform X2; mannan synthase 1-like isoform X1; polygalacturonase inhibitor; nuclear transcription factor Y subunit B-10; and serine/threonine protein receptor genes.

## Discussion

The use of polymorphisms embedded in *Fsp*-responsive DEGs successfully identified five QTLs. Dense linkage maps can be constructed due to the high density of SNPs found in the genomes and the high-volume data produced by RNAseq. This is an alternative approach to genotyping by sequencing. The identification of DEGs that respond to or are associated with specific biotic or abiotic stimulus, and the development of markers embedded in these gene sequences is an efficient approach for fine mapping and discovery of underlying genes for a desirable trait.

Here, we have reported the identification of five QTLs that are associated with *Fsp* resistance in pea (Table 3). Each of these QTLs explains 5.26-14.76% of the total phenotypic variation, and together they add up to 33.21% of the variation. To the best of our knowledge, this is the only study to report the presence of QTLs associated with *Fsp* resistance on chromosomes II and III of pea (Table 3).

A total of three QTLs were identified on chromosome V. Previously, three QTLs associated with *Fsp* resistance, *Fsp-Ps 3*.1, 3.2 and *3*.3, had been reported on chromosome V (Coyne et al., 2015, 2019). QTLs *Fsp-Ps 3*.2 and *Fsp-Ps 3*.3 are located close to each other. In fact, their LOD support intervals overlap (Coyne et al., 2015). These two QTLs are located adjacent to the newly identified *WB-Fsp-Ps* 3.1; however, their LOD intervals do not overlap. Further fine mapping should be able to determine if these three QTLs are in fact three, two, or only one QTL. The situation is similar in the case of QTLs *WB-Fsp-Ps* 5.1 (Table 3) and *Fsp-Ps 3*.1 (Coyne et al., 2015). LOD intervals from these two QTLs do not overlap either, although they are located close to one another on chromosome V. This proximity can mean that they represent one QTL.

Interestingly, this study did not find any QTLs on chromosome VI. *Fsp-Ps* 2.1 (Coyne et al., 2019) and a QTL identified by Feng et al., (2011) located on chromosome VI explained 44.4– 53.4% and 39% of the phenotypic variance, respectively. These two remain the major QTLs identified so far for *Fsp* resistance in pea. The absence of a major QTL on chromosome VI in this study could be due to the diversity of the parental source of resistance used in this study (005001 and 00-5007), versus what was used in previous studies (PI 180693, PI 557501, ‘Carman’) (Feng et al., 2011; Coyne et al., 2015, 2019).

Traits such as the root severity index, plant height, and plant weight did not show normal distribution in previous reports (Coyne et al., 2015, 2019). Furthermore, histogram analyses showed that the data distribution was skewed with an S-curve shape towards susceptibility or showed a bimodal distribution (Coyne et al., 2015, 2019). The large effect shown by *Fsp-Ps* 2.1 may explain the bimodal distribution seen in previous reports. Data presented in this study did not show a bimodal distribution but showed a trend towards normal distribution, which might explain the absence of QTLs with large effects.

We identified transgressive segregation in the two populations. Several of the transgressive lines showed enhanced resistance. For instance, a total of 29 and 5 RILs are more resistant than 00-5001 and 00-5007, respectively, based on their RDS scores. Genotypes 00-5001 and 00-5007 were previously characterized as high yielding varieties with important agronomics such as higher resistance to *Fsp*, resistance to Fusarium wilt, semi-leafless leaf type, and anti-lodging characteristics. Therefore, the transgressive lines with enhanced resistance might serve as potential candidate cultivars with good agronomics and even higher resistance to *Fsp*. Yields of these transgressive lines can be compared to elite cultivars under controlled and root rot conditions.

The molecular mechanisms underlying partial and total growth inhibition mediated by *Fsp* in a wide array of different pea breeding lines have been investigated and several genes have been proposed as potential candidates (Hadwiger, 2008, 2015; Williamson-Benavides et al., 2020). The establishment of associations between disease-related genes and tolerance, resistance, or susceptibility can facilitate the understanding of the possible mechanism(s) involved in the pathogenicity of *Fsp* in pea. *WB-Fsp-Ps* 5.1 was the major QTL identified in this study. Several candidate genes were identified within this QTL region. Among the DEGs identified in this QTL, an F-box/kelch-repeat protein (DN2516_c0_g1_i1) demonstrated reduced expression at 12 hr (FC = −24.4) after *Fsp* challenge in the partially resistant, but not in the susceptible genotypes. Nine F-box protein–coding genes have been found in the region of a highly dominant QTL that provides resistance to *A. euteiches*, a root rot pathogen in pea. (Djébali et al., 2009; Pilet-Nayel et al., 2009). F-box proteins are known to be involved in hormone regulation and in plant immunity (Guo and Ecker, 2003; Lechner et al., 2006). A nodulation signaling gene found in *WB-Fsp-Ps* 5.1 was also found to be upregulated at 0 hr (FC = 2.4) in susceptible genotypes when the expression levels were compared against the expression levels in partially resistant genotypes. It has been reported that a central regulator of symbiotic nodule development is determinant of susceptibility towards *A. euteiches* in *Medicago truncatula* (Rey et al., 2013). Interestingly, contig DN7423_c0_g1_i2 was identified as a hypothetical protein and the expression of this contig was significantly higher in the partially resistant genotypes than in the susceptible genotypes at 12 hours under control (FC=−50.2) and under *Fsp*-inoculated conditions (FC=−31.1). Auxin and ethylene responsive transcription factors did not show differential expression; however, they are also potential candidates driving the effect of *WB-Fsp-Ps* 5.1.

Several genes associated with disease resistance were located in *WB-Fsp-Ps* 5.2 QTL region (Table S10). Contigs DN19556_c0_g1_i1, DN2007_c0_g1_i4, DN77_c0_g1_i6 were identified as anthocyanin 5-aromatic acyltransferase, endochitinase PR4, and protein enhanced disease resistance 2. These genes showed differential expression; however, their expression was significantly higher in the susceptible genotypes compared to the partially resistant genotypes (Williamson-Benavides et al. 2020). The same pattern was observed for the TFs present in the *WB-Fsp-Ps 5.2*; NDR1/HIN1-like protein 10 and polygalacturonase inhibitors found in *WB-Fsp-Ps* 3.1; the RPM1-like and putative disease resistance protein RGA3 identified in *WB-Fsp-Ps* 2.1; as well as for a cluster of F-box proteins, a pathogenic type III effector avirulence factor, a pectinesterase/pectinesterase inhibitor, and a protein enhanced disease resistance 4-like gene found in *WB-Fsp-Ps* 5.3. This pattern of expression was also exhibited by contigs DN7670_c0_g1_i2 (located in *WB-Fsp-Ps* 5.1), DN175_c0_g4_i1 (*WB-Fsp-Ps* 5.2), DN261_c0_g1_i1 (*WB-Fsp-Ps* 5.2), DN8340_c0_g1_i2 (*WB-Fsp-Ps* 5.2), DN2650_c0_g3_i1 (*WB-Fsp-Ps* 2.1), which were identified as membrane receptors (Table S9, S10, S12). The high expression of any of these genes might be associated with disease susceptibility. However, further reverse genetics analyses will need to be performed to determine the dominant or recessive nature of these genes and QTL(s).

Cell death in the pea-*Fsp* interaction can help in the progression of *Fsp* infection due to the necrotrophic nature of the *Fsp* pathogen (Williamson-Benavides et al., 2020). The contig DN352_c0_g1_i17 located within *WB-Fsp-Ps* 5.3 was identified as CPR-5 protein which is known to negatively regulate the senescence and chlorotic lesions induced by pathogens when controlling programmed cell death (Bowling et al., 1997; Yoshida et al., 2002). This gene is highly suppressed in expression at 12 hr (FC = −3.03) after *Fsp* challenge in the susceptible genotype, which might trigger cell death.

Gene TRINITY_DN813_c0_g1_i4, identified as hypothetical protein L195_g026293 and located in *WB-Fsp-Ps* 2.1, was highly overexpressed in the partially resistant genotype when compared against the expression values in the susceptible genotypes under controlled conditions at 6 (FC=−4.76) and 12 hour (FC=−2.71) and under *Fsp*-inoculation at 0 (FC=−3.76) and 12 hours (FC=−2.29). BLAST search of contig TRINITY_DN813_c0_g1_i4 identified it as *Medicago truncatula* 1-aminocyclopropane-1-carboxylate oxidase homolog 1 (XM_003627980) (e-value: 5e-165, percentage identity: 84.5%). The 1-aminocyclopropane-1-carboxylate oxidase enzyme is involved in the production of ethylene. Jasmonate-induced defense responses, the expected response to counter the presence of necrotrophic pathogens such as *Fsp*, are known to be associated with elevation of 1-aminocyclopropane-1-carboxylate oxidase and also to increase the activity of defense-related enzymes and subsequent control of disease incidence (Yu et al., 2011; Dixit et al., 2016). Another R gene located in *WB-Fsp-Ps* 2.1 is the putative disease resistance protein RGA3; however, differential expression was not observed for this gene (Williamson-Benavides et al. 2020).

## Conclusions

The use of polymorphic DEGs for QTL mapping resulted in the identification of a new major QTL *WB.Fsp-Ps* 5.1. This outcome indicates that a combined gene expression and genetics approach seems effective in identifying genomic regions that may otherwise remain undetected especially for quantitative traits. It is to be noted that QTLs *WB.Fsp-Ps* 5.2, and 3.1 have large LOD support intervals necessitating fine mapping. Candidate genes, nested in each QTL will be instrumental in furthering our understanding of *Fsp*-pea interactions.

## Supporting information

Table S1. Primers, product amplicons

Table S2. 769 DEGs and SNPs

Table S3. Results 769 SNPs 8 Parents

Table S4. Screening Pop1

Table S5. Screening Pop2

Table S6. Position 769 SNPs in genome

Table S7. DatabaseForQTLAnalysisPop1

Table S8. DatabaseForQTLAnalysisPop2

Table S9_QTL5.1

Table S10_QTL5.2

Table S11_QTL5.3

Table S12_QTL2.1

Table S13_QTL3.1

## Author Contributions

AD, EB, RS, and BWB designed the study. BWB, GN, and EB performed the experiments and generated the data. BWB analyzed the data. LP provided the tolerant pea genotypes and *Fsp* isolates. AD supervised the study. All authors read and approved the final manuscript.

## Acknowledgements

The authors are grateful to Kurt Braunwart, CEO, ProGene Plant Research for critical discussions and his support for the project. The authors are grateful to Dr. Nnadozie Oraguzie for critical reading of the manuscript and their valuable suggestions. This manuscript has been submitted to bioRxiv as a preprint.

